# CBFβ-MYH11 interferes with megakaryocyte differentiation via modulating a gene program that includes GATA2 and KLF1

**DOI:** 10.1101/453845

**Authors:** Guoqiang Yi, Amit Mandoli, Laura Jussen, Esther Tijchon, Gaëlle Cordonnier, Marten Hansen, Bowon Kim, Luan N. Nguyen, Pascal Jansen, Michiel Vermeulen, Emile van den Akker, Jonathan Bond, Joost H.A. Martens

## Abstract

The inv(16) acute myeloid leukemia associated CBFβ-MYH11 fusion is proposed to block normal myeloid differentiation, but whether this subtype of leukemia cells is poised for a unique cell lineage remains unclear. Here, we surveyed the functional consequences of *CBFβ - MYH11* in primary inv(16) patient blasts, upon expression during hematopoietic differentiation *in vitro* and upon knockdown in cell lines by multi-omics profiling. Our results reveal that primary inv(16) AML cells share common transcriptomic signatures and epigenetic determiners with megakaryocytes and erythrocytes. Using *in vitro* differentiation systems, we reveal that CBFβ-MYH11 knockdown interferes with normal megakaryocyte maturation. Two pivotal regulators, GATA2 and KLF1, are identified to complementally occupy RUNX1 binding sites upon fusion protein knockdown, and overexpression of GATA2 partly restores megakaryocyte directed differentiation suppressed by CBFβ-MYH11. Together, our findings suggest that in inv(16) leukemia, the CBFβ-MYH11 fusion inhibits primed megakaryopoiesis by attenuating expression of GATA2/KLF1 and interfering with a balanced transcriptional program involving these two factors.

## Introduction

Core-binding transcription factors (CBFs) have been proposed to shape both stem cell self-renewal and differentiation, and their dysfunction could potentially lead to cancer pathogenesis^1^. The CBFs are heterodimeric complexes composed of two distinct subunits, alpha and beta^2^. The CBF α subunit is encoded by the RUNX family (usually RUNX1/AML1 in the hematopoietic cells) and directly contacts the DNA sequence, whereas the non-DNA-binding CBF β subunit is thought to facilitate stabilizing the DNA affinity of the CBF complex. CBFs are often mutated in acute myeloid leukemia (AML), for example in t(8;21) AMLs, characterized by expression of the *AML1-ETO* fusion gene, or inv(16) AMLs, delineated by the presence of the *CBFβ-MYH11* (CM) event^3^. *CBFβ-MYH11*, encodes a fusion protein between CBFβ and smooth muscle myosin heavy chain (SMMHC/MYH11), and is associated with AML FAB subtype M4Eo accounting for around 6% of AML cases^4-6^. However, our understanding of its roles in leukemogenesis remains incomplete.

Expression of CBFβ-MYH11 is able to disrupt normal myeloid differentiation, predispose for AML initiation, and cause full leukemia transformation upon the acquisition of additional genetic changes^7, 8^. A recent study revealed that CBFβ-MYH11 maintains inv(16) leukemia by obstructing RUNX1-mediated repression of MYC expression, which is featured by the replacement of SWI/SNF for PRC1 at MYC distal enhancers^9^. However, at which differentiation stage CBFβ-MYH11 blocks myeloid differentiation is still unclear. Mutational analysis of FACS-purified hematopoietic stem cells (HSC) as compared to leukemia cells confirmed the presence of CBFβ-MYH11 in HSCs, suggesting that the fusion event is involved in setting up a preleukemic cell state^10^. Further pursuing which differentiation pathway exactly is targeted by the oncoprotein would be needed.

At the molecular level, CBFβ-MYH11 in a complex with RUNX1 acts as a transcriptional regulator, which can depend on local genomic context, activate and repress genes involved in self-renewal, differentiation and ribosomal biogenesis^6, 11, 12^. Our previous findings have shown that a variety of cell surface markers increase in expression levels upon knockdown of CBFβ-MYH11 in the inv(16) cells, including those for the monocytic and megakaryocytic lineages^11^. In addition, mouse studies revealed that expression of the CBFβ-MYH11 protein causes abnormal erythropoiesis and gives rise to preleukemic pre-megakaryocyte/erythrocyte progenitors^8, 13^. Overall, these results potentially implicate a role of the CBFβ-MYH11 fusion in skewing cell differentiation orientation.

To investigate whether *CBFβ-MYH11* specifically blocks megakaryocyte/erythrocyte differentiation in the context of human hematopoiesis, and further probe its molecular mechanisms, we analyzed multiple transcriptomic and epigenomic profiles of inv(16) AMLs, several normal hematopoietic cell types and *in vitro* single-oncogene models. Our findings reveal a clustering of inv(16) AMLs towards megakaryocytes and erythrocytes based on DNA accessibility and H3K27ac-based super-enhancer profiles. Further molecular exploration indicates that CBFβ-MYH11 seems to be involved in interfering with normal differentiation through transcription deregulation and occupancy replacement of the transcription factors GATA2 and KLF1. Together, these results suggest that controlled expression of KLF1 and GATA2 expression is essential for inv(16) AML development.

## Materials and methods

### Human cells collection and sequencing

Leukemic samples were either obtained from bone marrow or peripheral blood for subsequent processing. Patients cells and cell lines were processed through multiple steps as previously reported^11^, and then subjected to high-throughput transcriptome and ChIP sequencing for histone marks, CBFβ-MYH11 fusion, RUNX1 and GATA2 as described in the Supplementary Information.

### Assays

Cell culture, differentiation of iPSCs towards the granulocytic lineage, nuclear extraction preparation, pulldown and Mass spectrometry were performed as detailed in the Supplementary Information.

### Bioinformatics analysis

#### Peak calling

After read mapping to the hg19 reference genome using BWA^14^ and removal of PCR duplicates by Picard *MarkDuplicates* option (http://broadinstitute.github.io/picard/), peak calling of CBFβ-MYH11 fusion, RUNX1, and GATA2 ChIP-seq was conducted using MACS1.3.3^15^ at a *p*-value cutoff of 10^-6^. For DNA accessibility and H3K27ac data in inv(16) patients, the detailed analyses procedure was described as before (Yi *et al*., submitted).

#### Super-enhancer identification

All H3K27ac peaks identified were filtered to exclude regions within +/- 2.0 kb around transcription start sites (TSSs), and then used as input for the ROSE algorithm^16^ to predict super-enhancers.

#### Tag counting

Read counts for each putative region were enumerated and then normalized to RPKM (reads per kilobase of gene length per million reads) for visualization in heat maps or boxplots. For each base pair in the genome, the number of overlapping sequence reads was determined, averaged over a 10 bp window and visualized in the UCSC genome browser (http://genome.ucsc.edu).

#### Motif analysis

Motif discovery was performed using GimmeMotifs^17^ with a threshold score of 0.9 (on a scale from 0 to 1).

#### Expression analysis

RNA-seq reads were uniquely mapped to the hg19 reference genome using STAR^18^, and subsequently normalized to RPKM values for all RefSeq genes using tag counting scripts. Gene-level count matrix was used as input for DESeq2 package^19^ to distinguish differentially expressed genes between any two groups. Significant genes were determined by a fold change cutoff of 1.5 and adjusted *p*-value of 0.01.

## Results

### CBFβ-MYH11 expressing cells harbor transcriptional characteristics of megakaryocytic and erythroid cells

To examine the transcriptional differences between inv(16) AML patients expressing CBFβ-MYH11 and normal hematopoietic cell types, we used RNA-seq and compared global gene expression levels between AML blasts and normal CD34^+^ progenitor cells (CD34), megakaryocytes (Mega), erythrocytes (Ery) and monocytes (Mono). Unsupervised principal component analysis (PCA) revealed that inv(16) AMLs displayed different transcriptomic landscapes as compared to normal lineages (Supplementary Figure 1A). The main source of variability (PC1) was the difference between Mega/Ery and CD34/inv(16) cells (Figure 1A), with the top contributing genes in PC1 mainly involved in immune response terms. In contrast, the second component (PC2) showed significant similarity between inv(16) and Mega/Ery, and was enriched in cell differentiation related terms (Figure 1A). The transcriptional difference between inv(16) leukemic blasts and normal cells was revealed by PC3, which terms associated with leukocyte migration and activation. Each pairwise comparison between individual normal cell type and inv(16) blasts revealed more than 2,000 differentially expressed genes (Figure 1B, and Supplementary Table 1). Together, these results suggest that inv(16) AMLs carry unique gene expression signatures (PC3), but also signatures that resemble CD34^+^ progenitor cells (PC1) or mature cells such as megakaryocytes (PC2).

**Figure 1.**
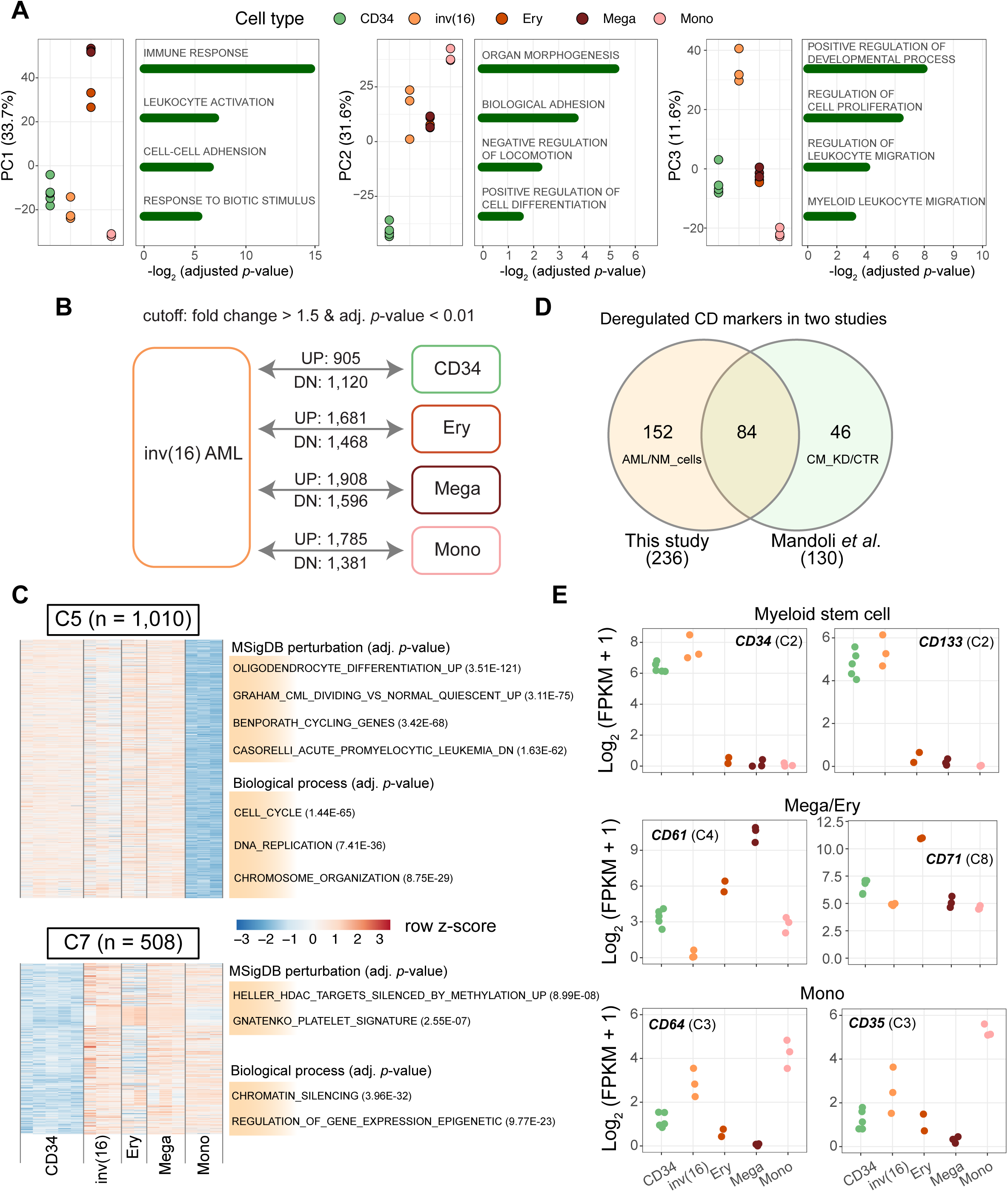
**The deregulated gene expression programs of primary inv(16) AML blasts compared to four normal cell types.** A. The transcriptional relationship based on RNA-seq among cell types revealed by principal component analysis. The five cell types are CD34: CD34^+^ progenitor cell, inv(16): primary inv(16) AML cell, Ery: erythrocyte, Mega: megakaryocyte and Mono: monocyte.
B. Pairwise gene expression comparison between primary inv(16) cells with other cell types.
C. Two example clusters after defining distinct expression patterns by k-means clustering among five cell types (see also Supplementary Figure 1B). The raw *p*-value in functional enrichment is adjusted by the Benjamini-Hochberg procedure.
D. Overlap of differentially expressed cell markers between our previous study (Mandoli *et al*., 2014) and the present study. NM_cells: normal cell types (CD34, Mega, Ery and Mono), CM_KD: CBFβ-MYH11 knockdown cells, CTR: control cells.
E. Transcriptional changes for several cell-type-specific markers. Labels in the parentheses indicate which cluster from Supplementary Figure 1B this gene is from.

Next, all differentially expressed genes were clustered into eight groups by the k-means clustering method (Supplementary Figure 1B, C). Besides cell-type-specific clusters (C1, C4, C6 and C8), we again found that inv(16) blasts shared gene signatures with normal cell types. For instance, the C2 cluster revealed similar expression patterns between inv(16) and CD34^+^ progenitors, while the inv(16) genes expressed in C3 resembled more closely a Mono signature. Furthermore, the primary inv(16) blasts shared a subset of gene signatures with Mega/Ery as shown in the C5 and C7 cluster (Figure 1C), although these genes also displayed similar expression levels in CD34 or Mono. These findings indicate that inv(16) blasts maintained certain transcriptional signatures from progenitors, but also already obtained characteristics of mature cells.

Different hematopoietic cell types can be identified by examining expression of cluster of differentiation (CD) markers. Here, we identified 236 CD markers differentially expressed between inv(16) and normal cells. Our previous studies showed altered transcriptional activity of CD markers upon transient knockdown of CBFβ-MYH11 in ME-1 cells^11^. To examine whether the 236 differentially expressed CD markers from this study might be directly regulated by CBFβ-MYH11 we compared the two datasets. More than half of CBFβ-MYH11 dependent CD markers (84/130) were also differentially expressed in the comparison of normal cell types versus inv(16) AMLs (Figure 1D), suggesting putative regulation by CBFβ-MYH11. Amongst these, myeloid stem cell markers like CD34 and CD133 displayed significantly higher expression levels in inv(16) AML than other mature cells (Figure 1E). However, some markers of mature cells like CD64 and CD71 were also expressed in inv(16) AML, suggesting inv(16) cells might already be partly differentiated.

To extend this finding we examined several other marker genes involved in hematopoiesis including GFI1B, RUNX1 and GATA2 (Supplementary Figure 2A). It has been shown that expression of *GFI1B* is key to megakaryocyte and platelet development^20^. Moreover, CBFβ-MYH11 binds a putative downstream regulatory element of *GFI1B* gene^11^ (Supplementary Figure 2B). We confirmed the highest transcriptional level of *GFI1B* in Ery/Mega, but also found that the inv(16) blasts expressed higher levels of *GFI1B* as compared to Mono, and similar to CD34 progenitors (Supplementary Figure 2A). Together, our results suggest that expression of CBFβ-MYH11 leads to impaired differentiation in part through deregulation of genes involved in maturation of megakaryocytic cells.

### Epigenomic clustering of inv(16) cells, megakaryocytes and erythroblasts

To further investigate whether cells blocked by CBFβ-MYH11 are poised for a certain lineage we compared epigenetic landscapes. For this, we generated DNaseI-seq and H3K27ac ChIP-seq profiles in inv(16) AML cells and compared those to profiles created in Mega, Ery and Mono. In addition, we downloaded public ATAC-seq data^21^, which showed high consistency with DNaseI-seq (Supplementary Figure 3A), from several progenitor cell types including hematopoietic stem cell (HSC), common myeloid progenitor (CMP), granulocyte-macrophage progenitor cell (GMP) and megakaryocyte-erythroid progenitor cell (MEP), to assess the similarity with inv(16) AMLs in open chromatin patterns. Principal component analysis (PCA), the t-distributed stochastic neighbor embedding (t-SNE) and hierarchical clustering results based on DNA accessibility showed robust classification of cell types and differentiation trajectory (Figure 2A, B). Interestingly, the inv(16) AML cells displayed most similarity in open chromatin signatures and hence closer relationships with Mega and Ery cells (Figure 2B, Supplementary Figure 3B).

**Figure 2.**
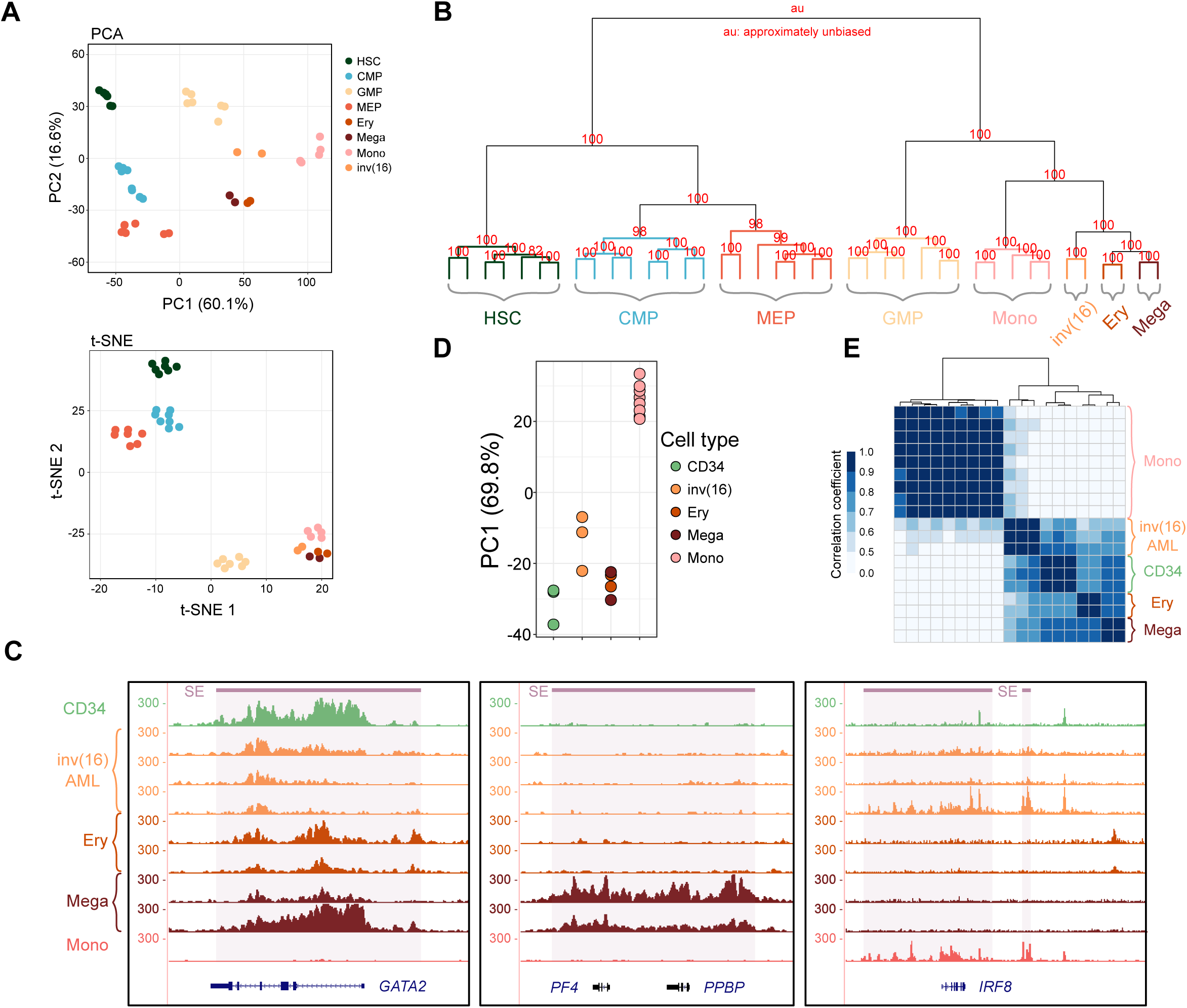
**Primary inv(16) cells are epigenetically more similar to megakaryocytes and erythrocytes.** (A) PCA and t-SNE analysis of DNA accessibility data display a clear separation along cell differentiation trajectories. Normal cell types include HSC: hematopoietic stem cell, CMP: common myeloid progenitor, GMP: granulocyte-macrophage progenitor cell, MEP: megakaryocyte-erythroid progenitor cell, Ery: erythrocyte, Mega: megakaryocyte and Mono: monocyte. inv(16): primary inv(16) AML cell.
(B) Hierarchical clustering of the top 5,000 variable DNA accessibility sites. Numbers on the branch indicate bootstrap support scores over 1,000 samplings.
(C) Variable super-enhancer landscapes (H3K27ac ChIP-seq) in the *GATA2*, *PF4*/*PPBP* and *IRF8* loci. SE: super-enhancer. Average H3K27ac density of three CD34 cells and nine monocytes were calculated for better visualization.
(D-E) Principal component 1 (PC1) and clustering plots of cell type relationship based on H3K27ac signal in super-enhancers.

Given that super-enhancers (SEs) are able to precisely capture cell identity^22^, we delved into genome-wide SEs landscapes using H3K27ac profiling. To integrate progenitor cells in our SE analysis, we downloaded H3K27ac data of CD34^+^ cells from NIH Roadmap Epigenomics^23^. The profiles of CD34, inv(16), Mega, Ery and Mono demonstrated high-quality H3K27ac binding patterns and cell-type-specific SEs profiles, for example at the *GATA2, PF4* and *IRF8* loci (Figure 2C, Supplementary Table 2). PCA analysis of H3K27ac signal at SEs confirmed these findings and showed that principal component 1 could clearly separate Mono from CD34, inv(16), Mega and Ery, and also reveal relatively closer distance between inv(16) and Mega/Ery (Figure 2D, Supplementary Figure 3C). Pearson clustering of super-enhancers again uncovered preferential grouping of inv(16) cells with CD34, Mega and Ery, while Mono super-enhancers formed an individual cluster (Figure 2E). Overall, these results suggest that consistent with our RNA-seq findings, inv(16) AML cells might carry a specific epigenetic state with high similarity to Mega/Ery cells, and represent cells that are putatively blocked along the Mega/Ery differentiation pathway.

### *CBFβ-MYH11* single oncogene expression blocks megakaryocyte/erythrocyte differentiation

A major limitation when analyzing inv(16) cell lines and primary inv(16) AML blasts is that they harbor many additional mutations. To exclude the disturbance of other genetic lesions, we utilized an *in vitro* iPSC-based hematopoietic differentiation system in which the expression of a single oncogene can be induced with doxycycline (dox), as successfully conducted in our previous study^24^. The established system contains dox-inducible CBFβ-MYH11 (Figure 3A, B) and we expressed the oncoprotein during differentiation towards the granulocytic, monocytic and megakaryocyte lineage (Figure 3C), allowing the investigation of the effects of CBFβ-MYH11 in the absence of additional leukemia driver mutations. As compared to cells in which CBFβ-MYH11 expression was not induced (control), two independent experiments revealed a remarkable reduction in CD41a, CD42b and CD235a positive cells during megakaryocyte differentiation (Figure 3D). In contrast, no effects were observed during monocyte and granulocyte/neutrophil differentiation. Together, these results reflect that CBFβ-MYH11 has a dominant effect on blocking Mega/Ery differentiation, and suggest inv(16) cells might correspond to cells that are arrested in a progenitor stage of this lineage.

**Figure 3.**
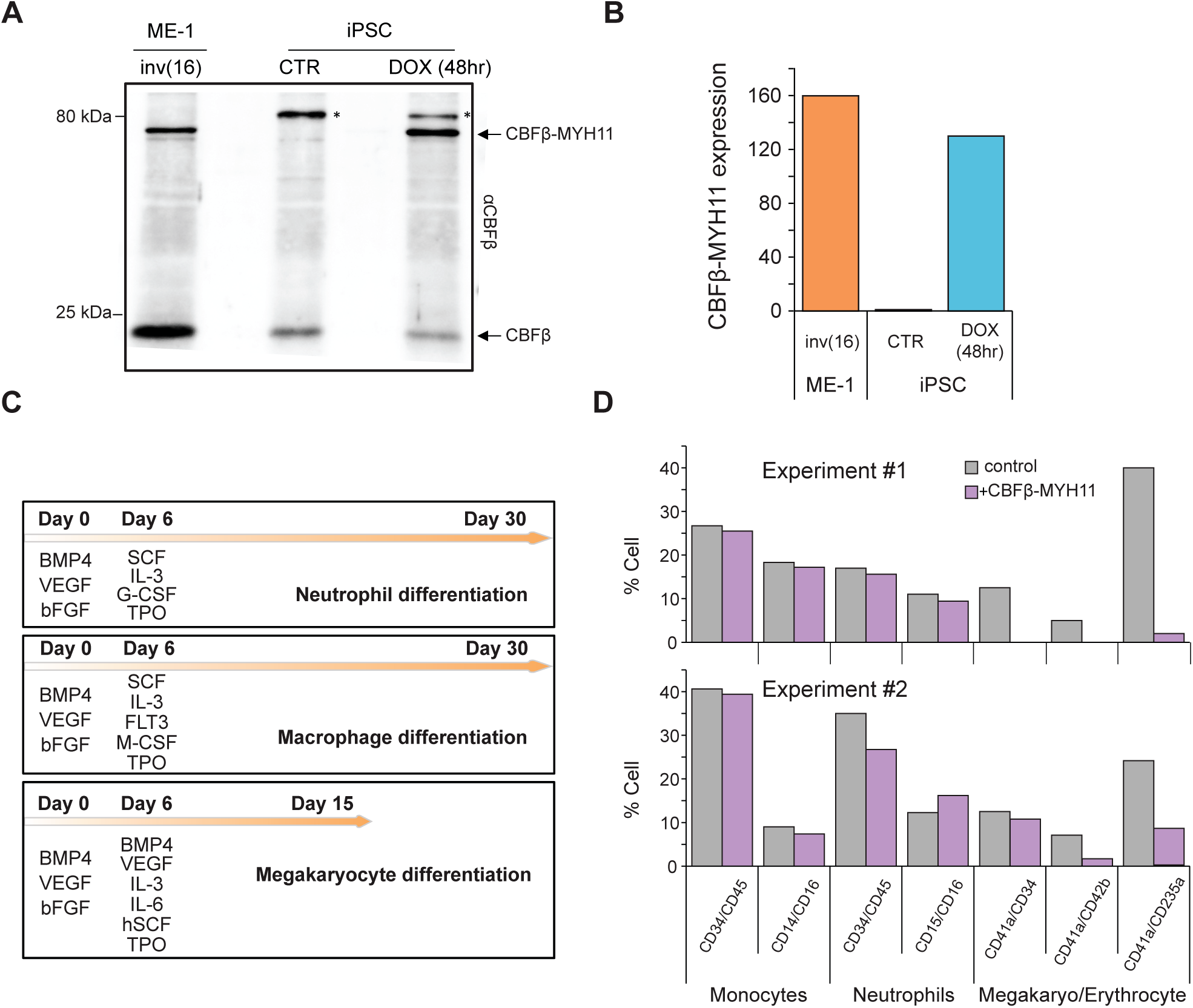
**The *CBFβ-MYH11* oncogene blocks megakaryocyte/erythrocyte differentiation.** A. Western blots of the CBFβ-MYH11 oncoprotein in the inv(16) AML ME-1 cell line and in the iPSC system before and after dox addition. Arrows point to the endogenous CBFβ and inducible expressed CBFβ-MYH11 protein. Asterisks represent aspecific binding of the antibody. Molecular weight markers in kilodaltons are listed on the left.
B. Expression levels of the *CBFβ-MYH11* oncogene in ME-1 cells and in iPSC without (CTR) or after dox induction of CBFβ-MYH11.
C. The methodology for iPSC differentiation towards neutrophil, macrophage and megakaryocyte lineages.
D. The percentage of cells showing specific myeloid markers after differentiation following the protocols in (C). Results for two independent differentiation experiments are shown.

### CBFβ-MYH11 knockdown affects cell proliferation

To further explore the detailed functional pathways induced by CBFβ-MYH11 alone, we generated a stable dox-inducible CBFβ-MYH11 knockdown (KD) cell line based on the inv(16) cell line ME-1^25^ (Figure 4A), and further examined genome-wide changes in the transcriptome as well as the acetylome. Previously, CBFβ-MYH11 transduction of CD34^+^ cells was shown to enhance proliferation^26^. In line with these results, knockdown of CBFβ-MYH11 in ME-1 cells affected proliferation, with more cells in G1-phase, while a slight reduction in cell viability was observed (Figure 4B, C). The latter finding was also supported by an increased number of cells in the sub-G1 cell phase, which is indicative of apoptotic or necrotic cell death (Figure 4C, Supplementary Figure 4). Transcriptome analysis between control and CBFβ-MYH11 knockdown cells at day 5 corroborated the effect on cell cycle, with genes involved in controlling DNA replication and cell cycle pathways increased in expression level (Figure 4D). Together these results suggest that CBFβ-MYH11 knockdown might trigger the onset of cell cycle arrest^27^.

**Figure 4.**
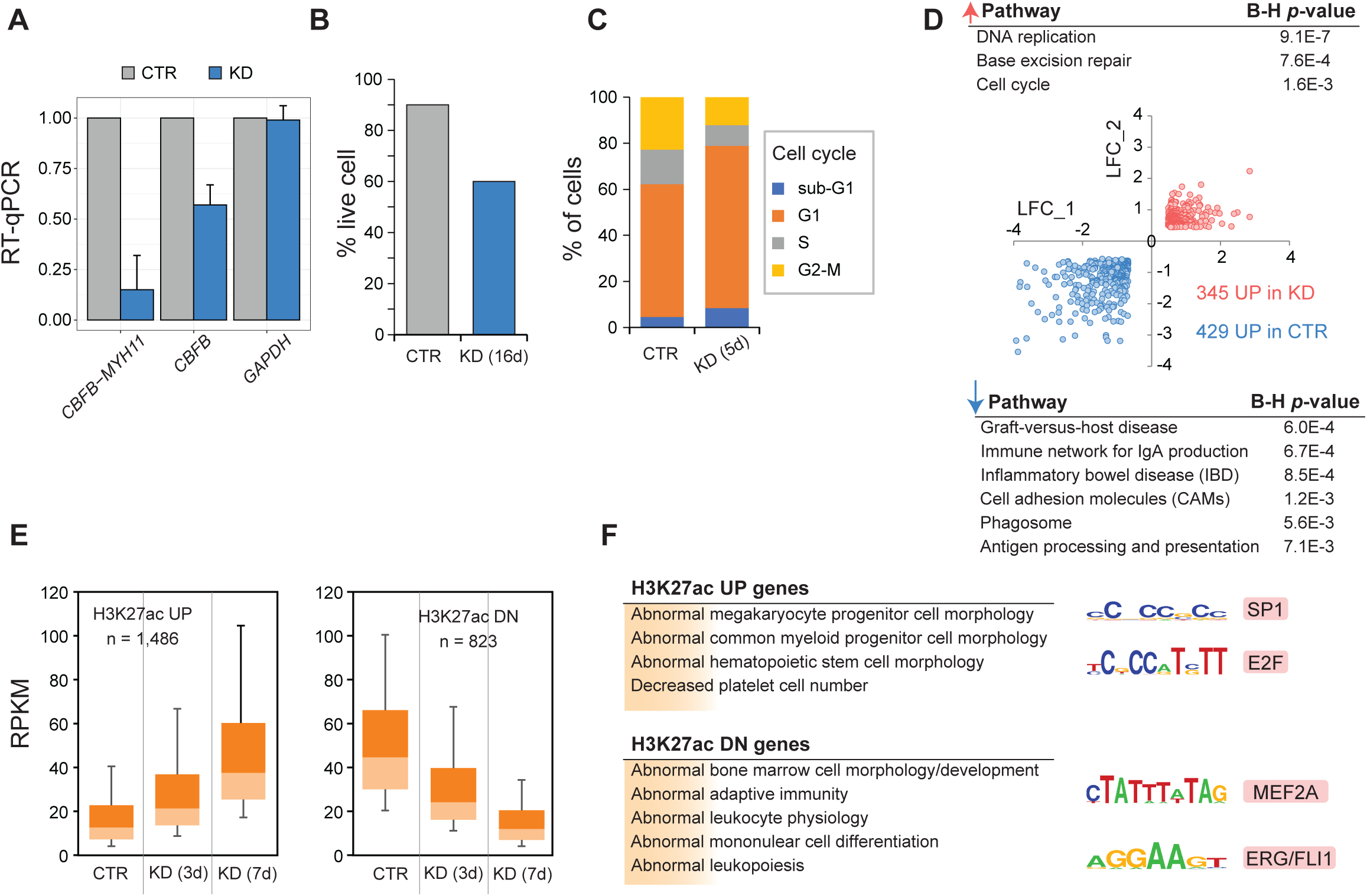
**Cell cycle deregulation induced by CBFβ-MYH11 fusion.** A. RT-qPCR analysis of *CBFβ-MYH11* and *CBFβ* in ME-1 cells before and after CBFβ-MYH11 knockdown (KD). Data are normalized to *GAPDH* expression level.
B. Cell viability in response to CBFβ-MYH11 knockdown.
C. FACS analysis showing the percentage of cells in four cell cycle phases after CBFβ-MYH11 knockdown.
D. Log_2_ fold change (LFC) of ME-1 CBFβ-MYH11 knockdown vs ME-1 control cells for a replicate RNA-seq experiment. Involved pathways of differentially expressed genes between control and CBFβ-MYH11-knockdown cells are indicated.
E. Differential H3K27ac enrichment before (CTR) and after (KD) CBFβ-MYH11 knockdown.
F. Functional annotation of genes and enriched motifs associated with differential H3K27ac peaks.

Given that altered transcriptional levels are a reflection of epigenetic changes, we performed H3K27ac ChIP-seq (positively correlated with gene expression) on control and CBFβ-MYH11 knockdown cells, and inspected differential H3K27ac peaks (> 100 tags and 2-fold difference). A total of 2,309 differential regions were identified, around 16.2% of them were covered by super-enhancers from inv(16) AML patients. The genes associated with regions showing increased acetylation after knockdown were functionally related to megakaryocyte differentiation and platelet formation (Figure 4E, F), while genes assigned by loci going down in acetylation were involved in leukocyte physiology. Motif analysis in the two types of peaks revealed enrichment for MEF2A and ERG/FLI1 motifs in regions attenuated in H3K27ac signal (Figure 4F, right). In contrast, H3K27ac-increased regions were significantly occupied by SP1 and E2F motifs, again suggesting deregulation of the DNA replication machinery^28^.

### CBFβ-MYH11 knockdown increases GATA2/KLF1 occupancy at RUNX1 binding sites

As we observed significant changes in transcription and H3K27ac occupancy after CBFβ-MYH11 knockdown, we set out to further excavate its regulatory mechanisms. As CBFβ-MYH11 binds the RUNX1 protein, it potentially interferes with the normal RUNX1-regulated gene program. To investigate the proteins which might be implicated in altering the transcriptional program, we performed DNA pull-down experiments using a specific nucleotide sequence bound by CBFβ-MYH11 that contains the RUNX1 core consensus motif TGTGGT (RUNX1 oligo), in control and CBFβ-MYH11-knockdown ME-1 cells (Figure 4A). We could show that the oligonucleotide with the RUNX1 motif efficiently pulled down CBFβ-MYH11 as well as RUNX1 and CBFβ from an ME-1 cell lysate (Figure 5A), whereas significantly attenuated affinity for CBFβ-MYH11 was observed when the knockdown cell lysate was used. Importantly, knockdown of CBFβ-MYH11 did not affect RUNX1 and wild-type CBFβ occupancy, suggesting changes in protein binding profiles are likely due to the absence of CBFβ-MYH11.

**Figure 5.**
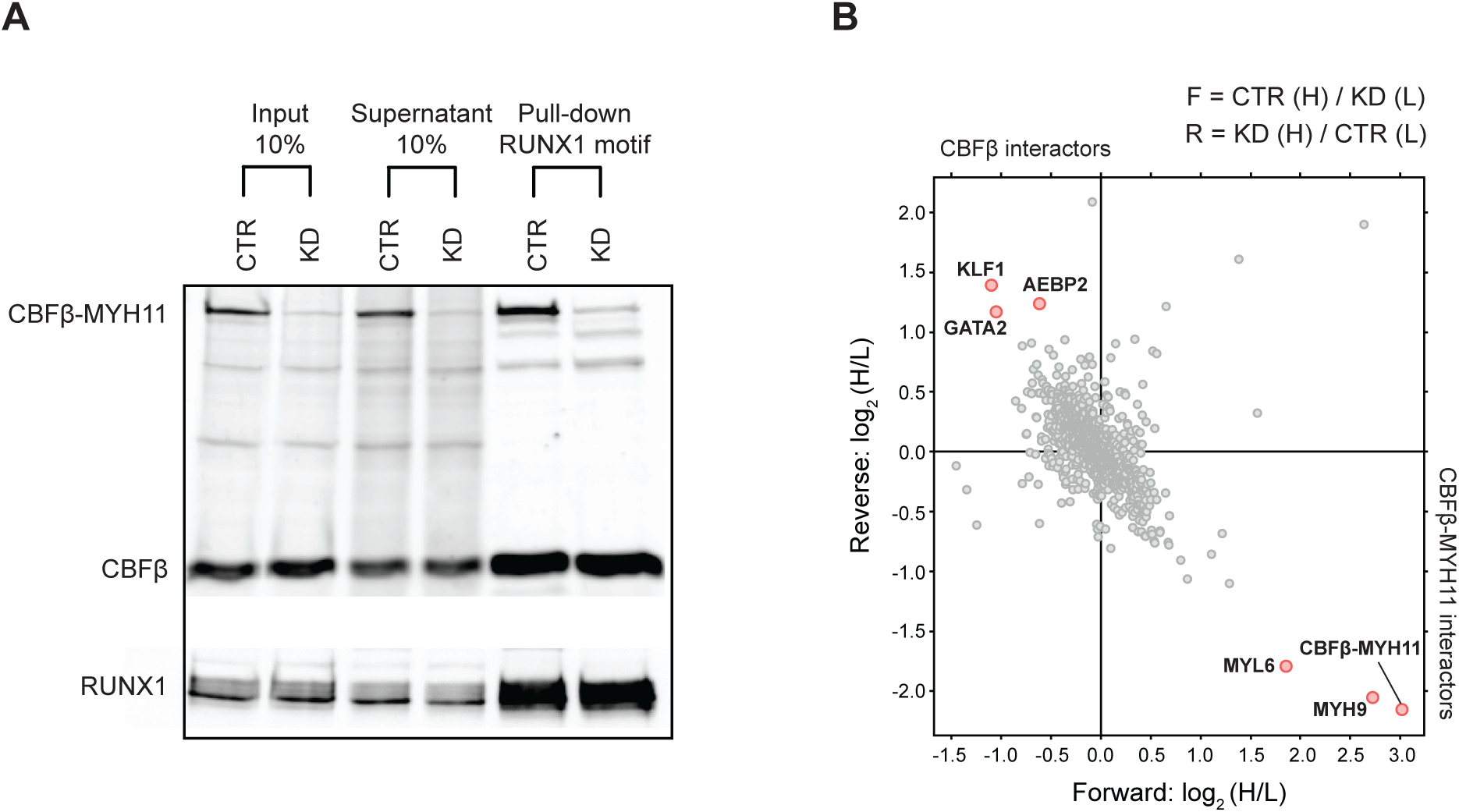
**Identification of interactors affected by the CBFβ-MYH11 fusion.** A) Western blot analysis of a DNA pull-down experiment in ME-1 cells using CBFβ and RUNX1 antibodies. B) Scatterplot showing the CBFβ-MYH11 interactome. Proteins interacting with a RUNX1 motif containing oligo before or after CBFβ-MYH11 knockdown are plotted by their SILAC-ratios in the forward (x-axis) and reverse (y-axis) SILAC experiment. Specific interactors with the RUNX1 motif containing oligo that bind upon CBFβ-MYH11 protein depletion lie close to the diagonal in the upper left quadrant. Interactors for which binding is CBFβ-MYH11 dependent are in the lower right quadrant.

To decipher the interactome of the CBFβ-MYH11 complex at RUNX1 binding sites, we employed previously described SILAC-based technology^11, 29^, using extracts derived from CBFβ-MYH11-knockdown ME-1 cells grown in light (L) or heavy (H)-labelled medium, incubated with oligonucleotides containing the RUNX1 motif (see Methods section). Of the > 1,300 identified proteins at a high confidence level, only a limited number displayed highly significant SILAC-ratios (>2) as CBFβ-MYH11 interactors. The CBFβ-MYH11 complex seems to facilitate the recruitment of MYH9 and MYL6 proteins to RUNX1 sites, as shown by their enrichment in the pull-down assay from ME-1 lysates (Figure 5B). Interestingly, upon CBFβ-MYH11 knockdown, two transcription factors (TFs), GATA2 and KLF1, are strongly enriched at RUNX1 occupancy loci, implying that these might play a crucial role in coordinating the RUNX1-dependent regulatory network in normal cells. Our previous findings demonstrated that *GATA2* and *KLF1* displayed enhanced levels in transcription after CBFβ-MYH11 knockdown^11^, and the present work could confirm not only increased RNA expression, but also stronger binding to the RUNX1 containing oligo by western blot (Supplementary Figure 5A, B), suggesting that the two TFs are repressed and might be replaced by CBFβ-MYH11 fusion in inv(16) AML. These findings are further supported by our previous inability to detect GATA2 and KLF1 as binders of RUNX1 sites^11^, as these assays were also done in the presence of CBFβ-MYH11.

### GATA2/CBFβ-MYH11 switching drives megakaryocyte/erythrocyte differentiation

As GATA2 is the highest expressed GATA factor in ME-1 cells, we investigated the genome-wide binding pattern by ChIP-seq in control and CBFβ-MYH11-knockdown inv(16) cells, and also examined the RUNX1 binding profile in parallel. The genome-wide screen indicated that GATA2 occupied similar genomic regions as RUNX1 at most locations. Quantifying GATA2 occupancy at RUNX1 loci revealed 2,410 sites with increased GATA2 signal and only 25 reduced binding sites. The 2,410 regions showed increased RUNX1 occupancy but reduced CBFβ-MYH11 binding (Figure 6A), and associated genes were involved in megakaryocyte/erythrocyte differentiation and platelet formation. Furthermore, genes associated with the 2,410 regions were increased in expression levels, which was further reflected by elevated occupancy level of the H3K27ac mark (Figure 6B, C), indicating activation of the GATA2-target gene program after the knockdown of CBFβ-MYH11. As could be expected, motif discovery at these regions presented remarkable enrichment for the GATA motif (Figure 6D). In addition, also overrepresentation for the EGR motif was detected, suggesting that TFs binding this motif are potentially involved in modulating the gene transcription process in a synergistic manner (Figure 6D).

**Figure 6.**
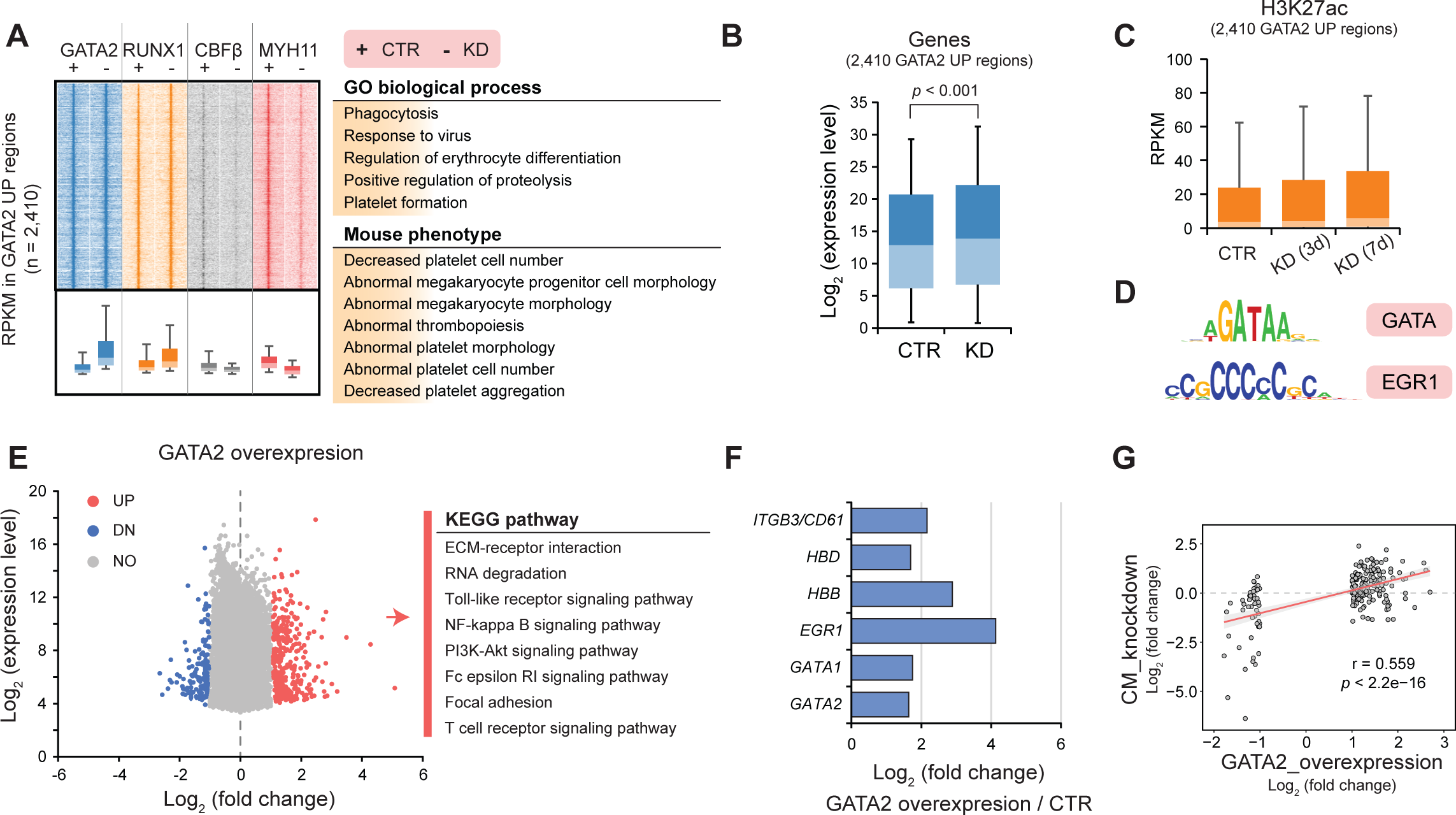
**Overexpression of GATA2 can partly restore cell differentiation towards megakaryocytes.** (A) Associated regulators and functional enrichment (ChIP-seq) in regions with increased GATA2 intensity after CBFβ-MYH11 knockdown (KD).
(B-C) Genomic loci with elevated GATA2 signal show higher expression levels (B) of associated genes and H3K27ac occupancy (C) in the CBFβ-MYH11 knockdown cells.
(D) Motif enrichment in regions with increased GATA2 density.
(E) Gene changes upon GATA2 expression in ME-1 cells. (right) Enriched KEGG pathways of differentially expressed genes after enforced GATA2 expression.
(F) Upregulated cell marker genes induced by GATA2 overexpression.
(G) Correlation of deregulated genes between ME-1 cells overexpressing GATA2 and ME-1 cells upon CBFβ-MYH11 knockdown. Pearson correlation coefficient and *p*-value are shown.

To probe whether enforced GATA2 expression is sufficient to switch on this gene program, we transduced ME-1 cells with a GATA2 expression construct and performed global RNA-seq analysis. During hematopoiesis, overexpression of GATA2 shows a decrease in cell proliferation and induces differentiation toward the megakaryocyte lineage^30^. Here, global transcriptional analysis identified a total of 367 genes increased and 171 genes decreased in transcriptional levels after GATA2 overexpression (Figure 6E). As expected, these upregulated genes contained GATA2 and several specific marker genes linked to megakaryocytic/erythrocyte differentiation, such as *CD61*, *HBD*, *HBB*, *EGR1* and *GATA1* (Figure 6F), and were mainly associated with various signalling pathways. Comparing transcriptomic changes after GATA2 overexpression with those observed after CBFβ-MYH11 knockdown displayed moderate correlation (r = 0.559) based on relative expression ratio, and only showed the overlap of 20 genes (5.4%) increased in expression and 19 genes (11.1%) decreased. These results suggested that enforcement of GATA2 expression alone can partly restore cell differentiation towards megakaryocytes, despite insufficiency to completely mimic CBFβ-MYH11 knockdown.

Together, these findings show that activation of the GATA2-involved regulatory program after CBFβ-MYH11 knockdown is central but not sufficient in driving further differentiation, which is potentially orchestrated in collaboration with other factors like KLF1 and/or EGR1.

## Discussion

The *CBFβ-MYH11* fusion arising from inv(16) rearrangement was reported to lead to differentiation blockage of normal myeloid cells and result in AML, but whether this fusion blocks specific cell commitment has not been discerned. Here, we compared the global landscapes of gene expression, DNA accessibility and H3K27ac between primary inv(16) AML blasts and normal cell types, and further investigated the functional contributions of CBFβ-MYH11 and its involved regulatory program *in vitro*.

Comprehensive transcriptomic exploration *in vivo* not only consolidated our previous findings^11, 12, 31^, but pinpointed towards unique gene expression programs underlying inv(16) AML cells. Further epigenomic analyses indicated more shared epigenetic determiners between inv(16) AML blasts and Mega/Ery cells, suggesting that primary inv(16) AML cells might be epigenetically primed for Mega/Ery differentiation with some key transcriptional pathways blocked. Therefore, in inv(16) leukemogenesis, the CBFβ-MYH11 fusion might skew leukemic blasts towards the Mega/Ery phenotype by epigenetic predisposition^32^ or be involved in setting up a specific Mega/Ery differentiation block by altering the cells gene program.

We have shown previously that CBFβ-MYH11 could activate transcription of self-renewal genes, and also repress expression of differentiation markers in the context of the cell line model ME-1^11^. Here, we used *in vitro* models with a single mutation (expression of CBFβ-MYH11) for surveying the role of CBFβ-MYH11. Using an iPSC system with dox-inducible CBFβ-MYH11 revealed that this fusion is able to specifically block *in vitro* Mega/Ery differentiation, but hardly has any effects on granulocyte and monocyte maturation. This finding suggests expression of CBFβ-MYH11 affects specific cell differentiation pathways.

Previous studies have proposed that CBFβ-MYH11 impairs normal binding of other proteins due to the higher affinity with RUNX1, and alters expression of RUNX1-dependent target gene sets^11, 12, 33^. We found that *GATA2* and *KLF1* showed elevated expression levels^11^ and also acted as significant interactors of RUNX1 after CBFβ-MYH11 knockdown, suggesting that CBFβ-MYH11 alters normal transcriptional programs of the two regulators and competitively takes over their binding sites at RUNX1 occupancy loci. GATA2 and KLF1 exert key effects in the Mega/Ery lineage fate decision via interaction with other modulators^34-36^. Our RNA-seq analysis revealed attenuated expression levels of *GATA2* and *KLF1* in inv(16) blasts as compared to Mega/Ery, and that *GATA2* overexpression could partly restore megakaryocyte differentiation, suggesting that weak expression of *GATA2* is essential for inv(16) leukemia but that increased levels are needed for differentiation. Focusing on GATA2, ChIP-seq illuminated its transcriptional activation role at these RUNX1-dependent target genes by increased recruitment of histone acetyltransferase activity and putative other TFs like EGR1^37-39^. Consequently, the data suggests that the GATA2-involved binding/regulatory program might be obstructed by the CBFβ-MYH11 fusion, leading to block of differentiation towards Mega/Ery in inv(16) leukemogenesis.

Megakaryocyte and erythrocyte share plenty of common molecular signatures, but the two cell types also exhibit distinct patterns in subtle regulatory networks^34, 40^. High transcription of *GATA2* has been reported to solely promote megakaryocyte differentiation and suppress erythroid maturation^41^. In our study, GATA2 overexpression boosted expression levels of some typical marker genes driving Mega differentiation but did not display a strong correlation with results from CBFβ-MYH11 knockdown. This finding reveals that CBFβ-MYH11 interferes with coordinated orchestration of a precise balance of multiple factors in maturing Mega/Ery, but the enforced expression of GATA2 only recapitulates megakaryocyte differentiation during bifurcation of the two lineages. As a consequence, GATA2 alone is not sufficient to inhibit CBFβ-MYH11-caused leukemia, and it maybe has greater functional relevance only in context with overexpression of other regulators like KLF1.

In summary, our study suggests that the CBFβ-MYH11 fusion maintains inv(16) AML cells by attenuating expression levels of GATA2 and blocks their further differentiation towards Mega/Ery lineages via interfering with a GATA2/KLF1-involved regulatory network. Collectively, these results corroborate our previous findings and facilitate a better understanding in the role of CBFβ-MYH11 in leukemia.

## Author contributions

The work presented here was carried out in collaboration between all authors. J.H.A.M., G.Y., A.M., L.J., E.T., G.C., M.H., B.K., L.N.N., P.J., M.V., E.v.d.A. and J.B. designed methods and performed the experiments. J.H.A.M., G.Y., A.M. and L.J. interpreted the results and wrote the manuscript, and all authors contributed to the manuscript preparation. All authors reviewed and approved the final manuscript.

## Acknowledgments

We thank all patients for their sample donations used in this study. We are grateful to Dr. Bert van der Reijden for advice and discussion. This work was supported by the BLUEPRINT project (European Union’s Seventh Framework Programme grant agreement number 282510), the Dutch Cancer Foundation (KWF KUN 2009-4527 and KUN 2011-4937) and KIKA (project 311).

## Competing interests

The authors declare no competing financial interests.

---

**Supplementary Figure 1. Global transcriptional difference between inv(16) blasts and four normal cell types.**

A. Unsupervised principal component analysis in transcriptomics for five cell types.
B. Eight expression patterns by k-means clustering among five cell types and enriched functional terms for each cluster. The raw *p*-value in functional enrichment is adjusted by the Benjamini-Hochberg procedure.
C. Dynamic expression profiles for the eight clusters identified in (B) among five cell types.

**Supplementary Figure 2. Cell-type-specific changes for seven representative genes essential for normal hematopoiesis.**

A. Transcriptional levels in inv(16) AML cells and normal cell types for *GFI1B, ERG, GATA2, JAG1, KLF1, RUNX1* and *TAL1*. Labels in the parentheses indicate to which cluster this gene belongs.
B. UCSC track showing CBFB-MYH11 binding sites at a downstream region of the *GFI1B* gene. The ChIP-seq data for CBFβ and MYH11 was from ME-1 cells before (CTR) and after CBFβ-MYH11 knockdown (KD) in our previous study (Mandoli *et al*., 2014).

**Supplementary Figure 3. The difference in chromatin accessibility landscape across cell types.**

A. Correlation between DNaseI-seq data and ATAC-seq data in monocytes.
B. Open chromatin signatures of primary human hematopoietic cells. Number above the heatmap indicate how many biological replicates we used to calculate average density for each cell type.
C. Unsupervised principal component analysis based on H3K27ac signal in super-enhancers for five cell types.

**Supplementary Figure 4. Cell cycle analysis showing the percentage of cells in sub-G1 after CBFβ-MYH11 knockdown.**

**Supplementary Figure 5. Increased levels in both transcription and translation of GATA2 and KLF1 after depletion of CBFβ-MYH11 fusion.**

A. Fold change in expression levels of *GATA2* and *KLF1* by RNA-seq after depletion of the CBFβ-MYH11 fusion in the inv(16) AML ME-1 cell line.
B. Western analysis of a DNA pull-down experiment in ME-1 cells using GATA2 and KLF1 antibodies.

**Supplementary Table 1. Pairwise transcriptomic comparison between primary inv(16) cells and four normal cell types.**

**Supplementary Table 2. Super-enhancer profiles identified from each cell types.**

## References

1. Appleford PJ, Woollard A. RUNX genes find a niche in stem cell biology. J Cell Biochem 2009 Sep 1; 108(1): 14-21.

2. Speck NA, Gilliland DG. Core-binding factors in haematopoiesis and leukaemia. Nat Rev Cancer 2002 Jul; 2(7): 502-513.

3. Martens JH, Stunnenberg HG. The molecular signature of oncofusion proteins in acute myeloid leukemia. FEBS Lett 2010 Jun 18; 584(12): 2662-2669.

4. Cancer Genome Atlas Research Network. Genomic and epigenomic landscapes of adult de novo acute myeloid leukemia. N Engl J Med 2013 May 30; 368(22): 2059-2074.

5. Liu P, Tarle SA, Hajra A, Claxton DF, Marlton P, Freedman M, et al. Fusion between transcription factor CBF beta/PEBP2 beta and a myosin heavy chain in acute myeloid leukemia. Science 1993 Aug 20; 261(5124): 1041-1044.

6. Shigesada K, van de Sluis B, Liu PP. Mechanism of leukemogenesis by the inv(16) chimeric gene CBFB/PEBP2B-MYH11. Oncogene 2004 May 24; 23(24): 4297-4307.

7. Castilla LH, Garrett L, Adya N, Orlic D, Dutra A, Anderson S, et al. The fusion gene Cbfb-MYH11 blocks myeloid differentiation and predisposes mice to acute myelomonocytic leukaemia. Nat Genet 1999 Oct; 23(2): 144-146.

8. Cai Q, Jeannet R, Hua WK, Cook GJ, Zhang B, Qi J, et al. CBFbeta-SMMHC creates aberrant megakaryocyte-erythroid progenitors prone to leukemia initiation in mice. Blood 2016 Sep 15; 128(11): 1503-1515.

9. Pulikkan JA, Hegde M, Ahmad HM, Belaghzal H, Illendula A, Yu J, et al. CBFbeta-SMMHC Inhibition Triggers Apoptosis by Disrupting MYC Chromatin Dynamics in Acute Myeloid Leukemia. Cell 2018 Jun 28; 174(1): 172-186.e121.

10. Corces-Zimmerman MR, Hong WJ, Weissman IL, Medeiros BC, Majeti R. Preleukemic mutations in human acute myeloid leukemia affect epigenetic regulators and persist in remission. Proc Natl Acad Sci U S A 2014 Feb 18; 111(7): 2548-2553.

11. Mandoli A, Singh AA, Jansen PW, Wierenga AT, Riahi H, Franci G, et al. CBFB-MYH11/RUNX1 together with a compendium of hematopoietic regulators, chromatin modifiers and basal transcription factors occupies self-renewal genes in inv(16) acute myeloid leukemia. Leukemia 2014 Apr; 28(4): 770-778.

12. Cordonnier G, Mandoli A, Radhouane A, Hypolite G, Lhermitte L, Belhocine M, et al. CBFbeta-SMMHC regulates ribosomal gene transcription and alters ribosome biogenesis. Leukemia 2017 Jun; 31(6): 1443-1446.

13. Kuo YH, Landrette SF, Heilman SA, Perrat PN, Garrett L, Liu PP, et al. Cbf beta-SMMHC induces distinct abnormal myeloid progenitors able to develop acute myeloid leukemia. Cancer Cell 2006 Jan; 9(1): 57-68.

14. Li H, Durbin R. Fast and accurate short read alignment with Burrows-Wheeler transform. Bioinformatics 2009 Jul 15; 25(14): 1754-1760.

15. Zhang Y, Liu T, Meyer CA, Eeckhoute J, Johnson DS, Bernstein BE, et al. Model-based analysis of ChIP-Seq (MACS). Genome Biol 2008; 9(9): R137.

16. Whyte WA, Orlando DA, Hnisz D, Abraham BJ, Lin CY, Kagey MH, et al. Master transcription factors and mediator establish super-enhancers at key cell identity genes. Cell 2013 Apr 11; 153(2): 307-319.

17. van Heeringen SJ, Veenstra GJ. GimmeMotifs: a de novo motif prediction pipeline for ChIP-sequencing experiments. Bioinformatics 2011 Jan 15; 27(2): 270-271.

18. Dobin A, Davis CA, Schlesinger F, Drenkow J, Zaleski C, Jha S, et al. STAR: ultrafast universal RNA-seq aligner. Bioinformatics 2013 Jan 1; 29(1): 15-21.

19. Love MI, Huber W, Anders S. Moderated estimation of fold change and dispersion for RNA-seq data with DESeq2. Genome Biol 2014 Dec 5; 15(12): 550.

20. Monteferrario D, Bolar NA, Marneth AE, Hebeda KM, Bergevoet SM, Veenstra H, et al. A dominant-negative GFI1B mutation in the gray platelet syndrome. N Engl J Med 2014 Jan 16; 370(3): 245-253.

21. Corces MR, Buenrostro JD, Wu B, Greenside PG, Chan SM, Koenig JL, et al. Lineage-specific and single-cell chromatin accessibility charts human hematopoiesis and leukemia evolution. Nat Genet 2016 Oct; 48(10): 1193-1203.

22. Pott S, Lieb JD. What are super-enhancers? Nat Genet 2015 Jan; 47(1): 8-12.

23. Kundaje A, Meuleman W, Ernst J, Bilenky M, Yen A, Heravi-Moussavi A, et al. Integrative analysis of 111 reference human epigenomes. Nature 2015 Feb 19; 518(7539): 317-330.

24. Mandoli A, Singh AA, Prange KHM, Tijchon E, Oerlemans M, Dirks R, et al. The Hematopoietic Transcription Factors RUNX1 and ERG Prevent AML1-ETO Oncogene Overexpression and Onset of the Apoptosis Program in t(8;21) AMLs. Cell Rep 2016 Nov 15; 17(8): 2087-2100.

25. Yanagisawa K, Horiuchi T, Fujita S. Establishment and characterization of a new human leukemia cell line derived from M4E0. Blood 1991 Jul 15; 78(2): 451-457.

26. Wunderlich M, Krejci O, Wei J, Mulloy JC. Human CD34+ cells expressing the inv(16) fusion protein exhibit a myelomonocytic phenotype with greatly enhanced proliferative ability. Blood 2006 Sep 1; 108(5): 1690-1697.

27. Patel SR, Hartwig JH, Italiano JE, Jr. The biogenesis of platelets from megakaryocyte proplatelets. J Clin Invest 2005 Dec; 115(12): 3348-3354.

28. Cobrinik D. Pocket proteins and cell cycle control. Oncogene 2005 Apr 18; 24(17): 2796-2809.

29. Spruijt CG, Gnerlich F, Smits AH, Pfaffeneder T, Jansen PW, Bauer C, et al. Dynamic readers for 5-(hydroxy)methylcytosine and its oxidized derivatives. Cell 2013 Feb 28; 152(5): 1146-1159.

30. Koga S, Yamaguchi N, Abe T, Minegishi M, Tsuchiya S, Yamamoto M, et al. Cell-cycle-dependent oscillation of GATA2 expression in hematopoietic cells. Blood 2007 May 15; 109(10): 4200-4208.

31. Cai X, Gao L, Teng L, Ge J, Oo ZM, Kumar AR, et al. Runx1 Deficiency Decreases Ribosome Biogenesis and Confers Stress Resistance to Hematopoietic Stem and Progenitor Cells. Cell Stem Cell 2015 Aug 6; 17(2): 165-177.

32. Feinberg AP, Koldobskiy MA, Gondor A. Epigenetic modulators, modifiers and mediators in cancer aetiology and progression. Nat Rev Genet 2016 May; 17(5): 284-299.

33. Lukasik SM, Zhang L, Corpora T, Tomanicek S, Li Y, Kundu M, et al. Altered affinity of CBF beta-SMMHC for Runx1 explains its role in leukemogenesis. Nat Struct Biol 2002 Sep; 9(9): 674-679.

34. Dore LC, Crispino JD. Transcription factor networks in erythroid cell and megakaryocyte development. Blood 2011 Jul 14; 118(2): 231-239.

35. Kuvardina ON, Herglotz J, Kolodziej S, Kohrs N, Herkt S, Wojcik B, et al. RUNX1 represses the erythroid gene expression program during megakaryocytic differentiation. Blood 2015 Jun 4; 125(23): 3570-3579.

36. Tijssen MR, Cvejic A, Joshi A, Hannah RL, Ferreira R, Forrai A, et al. Genome-wide analysis of simultaneous GATA1/2, RUNX1, FLI1, and SCL binding in megakaryocytes identifies hematopoietic regulators. Dev Cell 2011 May 17; 20(5): 597-609.

37. Wu D, Sunkel B, Chen Z, Liu X, Ye Z, Li Q, et al. Three-tiered role of the pioneer factor GATA2 in promoting androgen-dependent gene expression in prostate cancer. Nucleic Acids Res 2014 Apr; 42(6): 3607-3622.

38. Hayakawa F, Towatari M, Ozawa Y, Tomita A, Privalsky ML, Saito H. Functional regulation of GATA-2 by acetylation. J Leukoc Biol 2004 Mar; 75(3): 529-540.

39. Wilson NK, Foster SD, Wang X, Knezevic K, Schutte J, Kaimakis P, et al. Combinatorial transcriptional control in blood stem/progenitor cells: genome-wide analysis of ten major transcriptional regulators. Cell Stem Cell 2010 Oct 8; 7(4): 532-544.

40. Psaila B, Barkas N, Iskander D, Roy A, Anderson S, Ashley N, et al. Single-cell profiling of human megakaryocyte-erythroid progenitors identifies distinct megakaryocyte and erythroid differentiation pathways. Genome Biol 2016 May 3; 17: 83.

41. Ikonomi P, Rivera CE, Riordan M, Washington G, Schechter AN, Noguchi CT. Overexpression of GATA-2 inhibits erythroid and promotes megakaryocyte differentiation. Exp Hematol 2000 Dec; 28(12): 1423-1431.

